# NLRP3 Inflammasome Exhibits Context-Dependent Roles in Glioblastoma-Astrocyte Crosstalk

**DOI:** 10.64898/2025.12.30.697027

**Authors:** Sushmita Rajkhowa, Lipika Sha, Durgesh Meena, Deepak Kumar, MS Revanth, Vikas Janu, Mayank Garg, Mohit Agrawal, Deepak Jha, Sushmita Jha

## Abstract

Glioblastoma (GBM) is a highly aggressive brain tumor characterized by molecular and cellular heterogeneity and poor patient survival. NLRP3 inflammasome regulates inflammation and cell death with context-specific roles in cancer, but its functions in GBM tumour and astrocyte interactions remain unclear. In this study, we analyzed NLRP3 expression and function in GBM cell lines (LN-229 and LN18), astrocytes (SVG), patient-derived tissues, and organoids. Basal NLRP3 expression was higher in astrocytes than in GBM cells and increased after LPS stimulation, altering astrocyte morphology and NLRP3 subcellular localization. siRNA-mediated silencing of NLRP3 reduced GBM proliferation, migration, and viability in GBM cells, while enhancing proliferation and cytokine secretion in astrocytes, highlighting its context-dependent effects. NLRP3 deficiency exhibited reciprocal cytokine paracrine signaling, with GBM cells releasing elevated levels of IL-6 and TNF-α, and astrocytes releasing elevated levels of IL-1β. Glyburide treatment reduced NLRP3 expression and IL-1β release in GBM cells but elevated NLRP3 and IL-1β in astrocytes. Overall, these data reveal NLRP3’s dual roles in GBM-astrocyte crosstalk, suggesting cell-type-selective inhibition for therapeutic exploration.

## Introduction

Glioblastoma (GBM) is the most aggressive primary brain tumor, characterized by extensive heterogeneity, diffuse invasiveness, and resistance to standard therapies(1–3). The GBM tumor microenvironment (TME) features heterogenous non-neoplastic cells, including 30–50% innate immune cells—predominantly microglia and macrophages—alongside cancer stem cells, astrocytes, and endothelial cells that collectively contribute to tumor maintenance and progression(4–6). Although NLRP3 is implicated in glioma pathogenesis its roles in GBM versus stromal astrocytes remain undefined. In GBM, glioma-derived cytokines, such as CX3CL1, recruit tumor-associated microglia and macrophages, supporting tumor growth and invasion (4). Elevated IL-1β within the tumor microenvironment enhances glioma proliferation, migration, and invasiveness (7). Astrocytes play a pivotal role in homeostasis as well as in response to inflammation and injury through coordinated signaling with microglia and endothelial cells (8). Conversely, astrocytes also provide metabolic and trophic support, secrete pro-tumorigenic cytokines, and remodel the microenvironment to favor GBM tumor invasion and immune evasion (9–11). Cytokine release that shapes a tumor-promoting microenvironment is controlled by innate immune sensing mechanisms, particularly the nucleotide-binding domain leucine-rich repeat-containing receptors (NLRs), which modulate inflammatory signaling and cell death through inflammasome formation and cytokine secretion (12,13).

Emerging evidence implicates the innate immune sensing pathways, especially NLRP3 inflammasome, as a key modulator of glioma pathogenesis (13–15). Upon activation, the NLRP3 inflammasome assembles into a multiprotein complex that facilitates caspase-1–dependent maturation of the pro-inflammatory cytokines IL-1β and IL-18, thereby driving inflammatory signaling and pyroptotic cell death (16). Although NLRP3 activation serves a protective role in host defense, dysregulated or persistent NLRP3 activity has been associated with chronic neuroinflammation and tumor-promoting processes within the central nervous system (17,18). While NLRP3 has been studied extensively for various diseases and cancer, very few reports have investigated GBM in the context of NLRP3 (13,16,17). Recent studies indicate that NLRP3 expression and inflammasome activity are upregulated in GBM tissues, particularly within the immunosuppressive tumor microenvironment, where NLRP3 signaling promotes the recruitment and expansion of granulocytic myeloid-derived suppressor cells (MDSCs) (19). Mechanistically, activation of the NLRP3 inflammasome in GBM promotes tumor growth and invasion by regulating inflammatory signaling cascades, stimulating cellular proliferation, and attenuating anti-tumor immune responses (14). These findings underscore the dual function of NLRP3 as both a sensor of cellular stress and a driver of tumorigenesis through modulation of the immune microenvironment. Consequently, therapeutic targeting of the NLRP3 inflammasome pathway represents a promising approach to enhance treatment efficacy and improve patient outcomes in GBM (15,19). In this study, we examine NLRP3 expression and activation across patient-derived glioma specimens and organoid models to interrogate the cellular sources of inflammasome activity in the GBM microenvironment and assess the impact of pharmacologic NLRP3 inhibition on tumor-intrinsic signaling and tumorigenic phenotypes. By integrating molecular profiling with functional perturbation, we aim to elucidate the role of NLRP3 in glioblastoma progression and assess its potential as a therapeutic target in precision oncology.

## Materials and Methods

### 1. Cell Culture

The human glioblastoma cell line LN-18 (RRID:CVCL_0392) was purchased from the American Type Culture Collection (ATCC), and the human glioblastoma cell line LN-229 (RRID:CVCL_0393) was purchased from the Cell Repository of the National Centre for Cell Science (NCCS) Pune, India. Human astrocyte cell line SVG was kindly shared by Dr Pankaj Seth from the National Brain Research Centre (NBRC), India. The astrocyte cells were analyzed using Western Blot with a GFAP antibody. Cells were grown in media containing DMEM (HiMedia, AL151A-500ML) along with 10% Fetal Bovine Serum (FBS) (Cell Clone, CCS-500-SA-U) and 1% antibiotic-antimycotic solution (penicillin, streptomycin, amphotericin B) (HiMedia, A002-200ML) at 37°C with 5% CO2.

### 2. PCR

RNA was isolated by the phenol-chloroform method, using TRIzol reagent (as per manufacturer’s instructions). The RNA concentration was quantified using a Nanodrop (ThermoScientific), and the quality was checked using both a UV-visible spectrophotometer (Nanodrop) and was used for first-strand synthesis PCR to prepare complementary DNA (cDNA) using iScript^TM^ cDNA synthesis kit (BIO-RAD 1708890). Endpoint PCR was performed using gene-specific primers for hypoxanthine-guanine phosphoribosyltransferase (HPRT-1), 18S, NLRP3, isocitrate dehydrogenase (IDH1, IDH2), epidermal growth factor receptor (EGFR), and tumor suppressor (TP53) to determine gene expression. SYBR green dye-based Real-time PCR (Agilent MX3000) was performed to quantify gene amplification and fold change in normal and glioma cells (New England BioLabs M3003S) using gene-specific primers. All results are normalized to HPRT, GAPDH and 18S ribosomal RNA internal controls and are expressed in relative numbers. Fold change was calculated using 2^△△^CT (20). A complete list of the primer and sequence details is provided in supplementary data (Table 1).

**Table 1.**
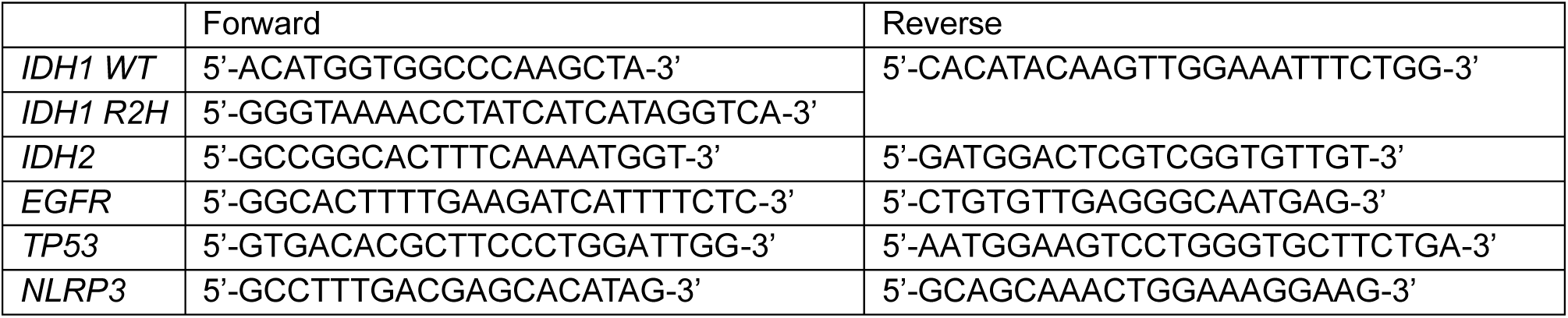
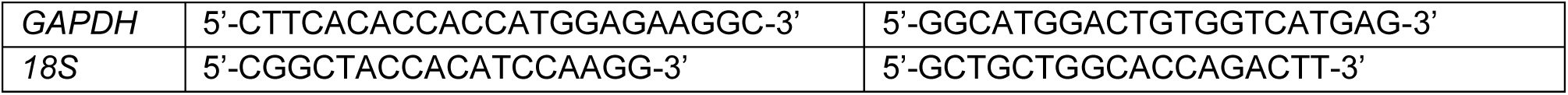
Primer Table list:

### 3. Immunocytochemistry

50,000 cells per well were seeded in DMEM (HiMedia, AL151A-500ML) along with 10% FBS (Cell Clone, CCS-500-SA-U) and 1% antibiotic-antimycotic solution (penicillin, streptomycin, amphotericin B) (HiMedia, A002-200ML) in chamber slides and incubated in a CO2 incubator (5% CO2, 37°C temperature, and 95% humidity). Cells were washed with PBS, fixed using 4% paraformaldehyde (HiMedia, MB059-500ML) for 10 minutes, permeabilized with 0.1% triton-X (Sigma, 1001723790) in PBS (permeabilization buffer) for 15 minutes, and blocked with blocking buffer containing 5% FBS in permeabilization buffer for 1 hour at 4°C (in a humidified chamber). Cells were immunolabelled with primary antibody (NLRP3: Novus Biologicals, NBP2-12446, actin: Santa-Cruz Biotechnology, mouse antibody, sc-47778, GFAP: Cell Signalling technology, mouse antibody, 3670S) and incubated overnight at 4°C. Subsequently, cells were washed with PBS and incubated with secondary antibody (Alexa fluor 488nm: Invitrogen, rabbit antibody, A11008; Alexa fluor 594nm: Invitrogen, mouse antibody, A11005) for 1 hour at 37°C (humidified and dark conditions), and slides were mounted with Vectashield containing DAPI (21). Immunopositive cells with an observable DAPI-stained nucleus were counted blindly twice.

### 4. Isolation of primary cells from patient tissues

Glioma tissues were obtained with approval from the Internal Review Board and the Ethics Committees of AIIMS, Jodhpur. Informed consent was acquired from human participants for the use of tissue samples for experiments. We have performed all experiments in accordance with the ethical guidelines and regulations of AIIMS, Jodhpur, and Indian Institute of Technology Jodhpur. Tissue sections were collected in aCSF (2mM CaCl2.2H2O (Calcium chloride dihydrate)), 10mM glucose, 3mM KCl (Potassium Chloride), 26mM NaHCO3 (Sodium bicarbonate), 2.5mM NaH2PO4 (sodium dihydrogen phosphate), 1mM MgCl2.6H2O (magnesium chloride hexahydrate), 20mM sucrose at AIIMS Jodhpur. Once received at our laboratory, aCSF was discarded, and the tissue was weighed. Tissues were washed in aCSF and PBS, chopped into approximately 1 mm-sized pieces, and transferred into a solution of 0.25% trypsin with EDTA (Ethylenediaminetetraacetic Acid). The tissue was dissociated by shaking at 250 RPM for 30 minutes at 37°C. After dissociation, neutralizing media were added to neutralize the trypsin, and the sample was centrifuged. The cell pellet is collected and resuspended in 1 mL of culture medium (45% DMEM, 45% F12 Nutrient medium, and 10% FBS). Cells were plated in a tissue culture flask and incubated in a CO2 incubator (5% CO2, 37°C temperature, and 95% humidity) (22).

### 5. Western blotting

Cells or tissues were lysed in radioimmunoprecipitation assay (RIPA) buffer with freshly added protease inhibitors for 4 min at 4°C. The protein concentration was assessed using a Bradford assay. 10µg of protein was loaded onto SDS-PAGE and then transferred onto a nitrocellulose membrane, which was incubated with blocking buffer (5% skimmed milk) for 1.5 hours, followed by primary antibodies (NLRP3: Novus Biologicals, NBP2-12446, β-actin: Santa-Cruz Biotechnology, mouse antibody, sc-47778) overnight and appropriate secondary HRP-conjugated antibodies (Anti-rabbit IgG HRP-linked: Cell signalling technology, 7074S, Anti-mouse IgG HRP-linked: Cell signalling technology, 7076S). Protein expression was visualized using the Azure Biosystems Gel Documentation system. ImageJ software was used for the densitometry analysis of the blots. The relative protein quantity was normalized to β-actin (23).

### 6. siRNA-mediated gene silencing

LN-229 cells were seeded in 6-well plates. After cell adherence, cells were left untreated (ctrl) or transfected with non-specific scrambled RNA control (scRNA 50nM) or *NLRP3* gene-specific siRNAs (50 and 100nM) in Opti-MEM medium. After 48 hours of treatment, transfected and control cells were lysed with RIPA buffer, and protein was isolated. Gene silencing was confirmed by western blot as described previously (21).

### 7. Wound healing assay

WT and *NLRP3*^-/-^ LN-229 cells were seeded in a 96-well plate (10,000 cells/well). After almost 90% confluency, a scratch was made in the well, and cells were observed for the next 24 hours (24). Images were taken using a multimode reader (cytation5, Agilent).

### 8. Colony formation assay

WT and *NLRP3*^-/-^ LN-229 cells were seeded at a density of 2500 cells per well (5% CO2; 37°C) in a chamber slide, and small colonies were observed after 36–48 hours. Colonies were stained with Giemsa, as previously described (21). Slides were observed using a bright field microscope, and results were quantified by counting the number of colonies formed per well and cells present per colony.

### 9. MTT

WT and *NLRP3*^-/-^ LN-229 cells were seeded at a density of 2500 cells per well (5% CO2; 37°C) in a 96-well plate. The next day, the media were removed from the wells. 90μL of serum-free DMEM media was added to the cells. MTT solution prepared in 1X PBS at a concentration of 12mM. 10uL of MTT solution to each well.The plate was placed in the incubator for 2 hours. After 2 hours, 100 μL of acidic isopropanol was added to each well, mixed properly, and kept for 10 minutes. Absorbance was taken at 570nM. For spheroids, 20μL of MTT solution was added to each well with 180μL serum-free DMEM media. The plate was placed in the incubator for 4 hours. After 4 hours, 200 μL of acidic isopropanol was added to each well, and the mixture was kept for 10 minutes after proper mixing. Absorbance was taken at 570nM.

### 10. Heterocellular Spheroid formation

WT and *NLRP3*^-/-^ LN-229 cells were seeded in low attachment U-shaped 96-well plate for spheroid generation. Cells were maintained in a medium containing 0.1% antibiotic,10% FBS, and nutrient media (DMEM) at 37 °C with 5% CO2. Cells began to aggregate within a few hours and formed spheroids within one day. The spheroids were imaged with a multimode reader (cytation5, Agilent) (23).

### 11. Glyburide treatment of cell lines

LN229 and SVG cells were seeded in a 6-well plate for 24 hours and then treated with 10 mM glyburide (dissolved in DMSO) for 30 minutes (Final Concentration 10μM) in serum-free nutrient media (DMEM) (25). The cells were incubated at 37 °C with 5% CO2. The supernatants were collected, and cells were seeded for monoculture and co-culture spheroids. Cells were then lysed with RIPA buffer, and protein was isolated.

### 12. Image acquisition of spheroid

The imaging of seeded spheroids was performed through an automated multimode reader (cytation5, Agilent) from day zero at 5% CO2 concentration and 37°C. Images were taken at different Z-planes, and after the imaging was completed, the Z-plane images were stitched using the built-in software. Final images were saved for further analysis.

### 13. Quantification of area, circularity, and compactness of spheroids

NIH Image J software was used to quantify the spheroid area (26). The spheroid area was manually outlined and measured using the application’s built-in tools. To assess the circularity and compactness of the spheroid, blinded observers were asked to score each category on a 5-point scale. In each category, the lack of a visible spheroid or formation of multiple spheroids was assigned a score of 1. For compactness, spheroids with clearly visible spaces and gaps were classified as a loose aggregate (score = 2), and aggregates with no gaps within the cell mass but presenting diffuse borders were scored as a tight aggregate (score = 3). Further compaction led to the formation of distinct dark borders around spheroids with few loose cells attached; these were classified as compact spheroids (score = 4). At the most compact stage, cells on the surface of the spheroid were remodelled and followed the contour of the spheroid, creating a smooth and defined outline, and were classified as tight spheroids (score = 5). For circularity, spheroids with similar degrees of concave and convex outline were classified as irregular (score = 2). In contrast, those mainly consisting of convex borders but with small concave dimples were classified as minor irregular (score = 3). Spheroids that are elongated with no concave outline sections receive a score of 4, and finally, symmetrically circular spheroids receive a score of 5 (Leung et al., 2015). Using the available data, a code was designed to automatically determine the area and score the circularity and compactness of the spheroids (13).

### 14. ELISA

Supernatants from cultured cells, patient-derived tissues, and gene knockdown heterocellular spheroids were analyzed for IL-1ß, IL-6, and TNF-α secretion by ELISA (BD Biosciences) as per manufacturer’s instructions.

### 15. Statistical Analyses

Unpaired Student’s t-tests were used to statistically evaluate significant differences. Data are presented as mean ± SEM. Differences were considered statistically significant if p<0.05.

## Results

### NLRP3 expression increases upon exposure to inflammatory signals

NLRP3, a pro-inflammatory protein, is one of the most broadly studied inflammasomes. Increased NLRP3 levels due to mutation have been reported to cause chronic auto-inflammatory diseases like Muckle Wells syndrome, familial cold-induced auto-inflammatory syndrome, metabolic diseases like diabetes, obesity, and cancer, like gastric cancer, hepatic cancer, colitis-associated cancer, etc. (13,21,27). To understand the expression of NLRP3 in GBM cell lines LN-229, LN-18, and astrocyte cells SVG, basal and LPS-induced cells were assessed. Quantitative PCR analysis revealed that LPS stimulation (0.5 ng/ul for h) induced a significant increase of >4 fold change in NLRP3 mRNA expression in both LN-229 and LN-18 cells as compared with untreated controls and normalized relative to GAPDH(n=3 ; p<0.05; Fig. 1C). Upon normalization to 18S housekeeping gene, >100 fold change in LN-229 cells and >1 fold change in LN-18 cells were observed (Fig. 1C). This upregulation was confirmed at the protein level by western blotting, supporting that inflammatory stimuli increase NLRP3 expression in tumor-derived cell lines (Fig. 1A). Immunocytochemistry revealed higher basal NLRP3 expression in LPS-stimulated astrocytes versus LN229, with LPS-induced punctate perinuclear localization and morphological changes in astrocytes (Fig. 1B, 1D; quantified blindly, n=3; >200 cells/field). These extend prior findings in patient-derived tissues, U87 and U251 cell lines, to new glioma cell lines and stromal cells (14). To our knowledge, this represents the first report of NLRP3 expression in the LN-229 and SVG cell lines. Together, these data establish that NLRP3 is present in both GBM cells and astrocytes and is regulated by inflammatory cues at the transcriptional and translational levels.

**Fig 1.**
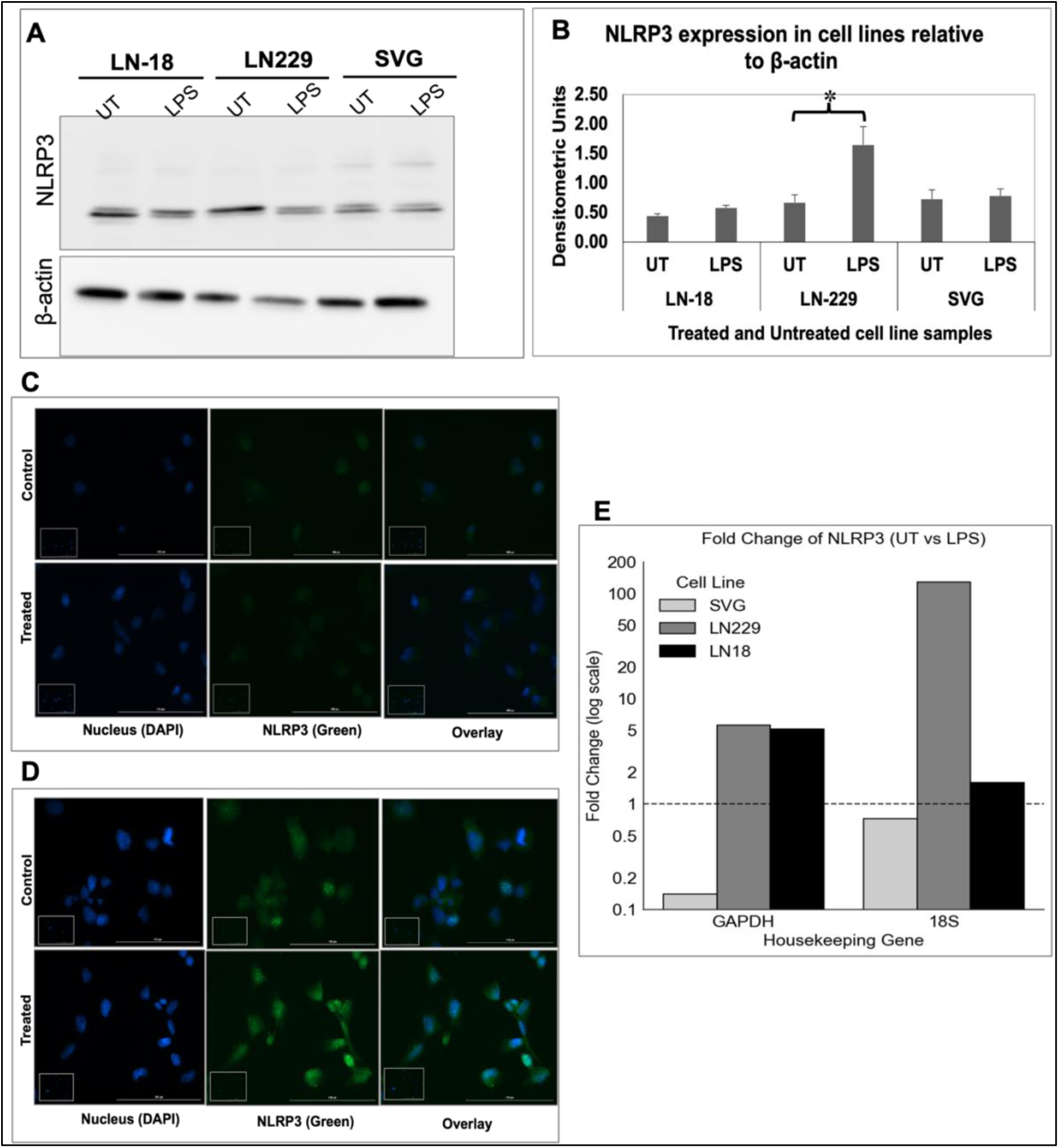
**NLRP3 expression in cell lines**: **A**. Western Blot for UT and LPS-treated GBM and astrocyte cells (n=3) **B**. Densitometric analysis normalized to beta-actin. **C.** Immunofluorescence of NLRP3 in GBM and **D**. astrocyte cells. **E**. Fold change of UT and LPS-treated GBM and astrocyte cells normalized to GAPDH and 18S genes (n=3). Data are represented as mean ± SD. Error bars indicated SD. ∗p < 0.05, **p < 0.01, ***p < 0.005 (Student’s t-test).

### *NLRP3* si-Knockdown Impairs Proliferation, Migration, and Viability in GBM Cells

To examine the functional significance of NLRP3 in GBM tumor biology, we performed siRNA-mediated knockdown of NLRP3 in LN-229 GBM and SVG astrocyte cell lines. Previous studies have reported that NLRP3 inhibition in Glioma cells, such as U87 and U251, suppresses cell proliferation, growth, and invasion (14). This demonstrates the tumor-promoting role of NLRP3 in glioma. Clonogenic assays revealed that NLRP3 depletion significantly reduced colony formation, migration, and cell viability in LN-229 GBM cells, indicating attenuated tumor growth (Fig. 2A-C). In contrast, NLRP3 knockdown increased all three parameters in SVG astrocyte cells (Fig. 2D- F), suggesting a cell-type-specific regulatory role.

**Fig 2.**
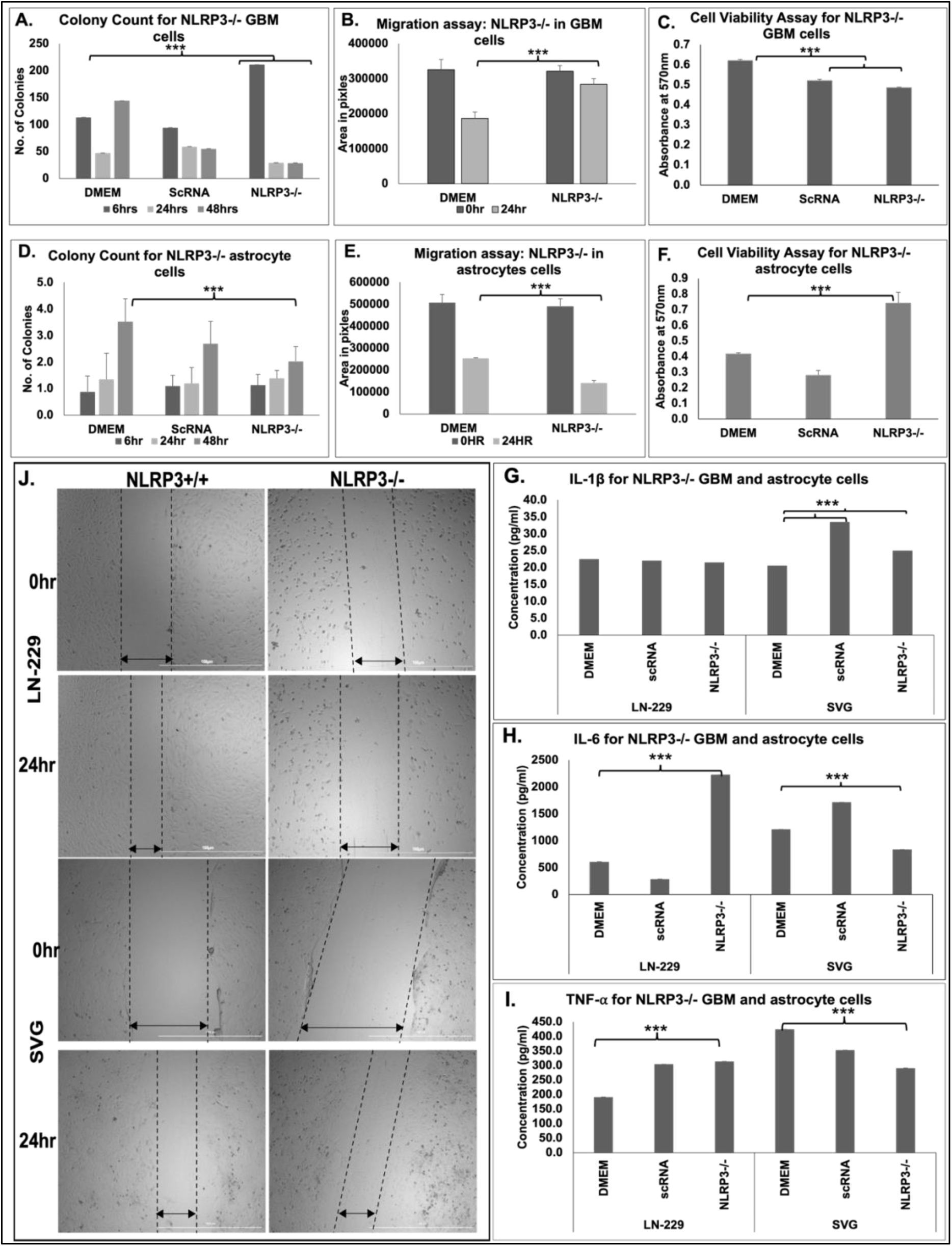
NLRP3 inhibition attenuates proliferation, migration, and viability in GBM cells. Quantification of CFA in **A**. GBM Cells. **D**. astrocyte cells. **J.** Wound healing Assay for *NLRP3-/*- in GBM and astrocyte cells. Quantification of migration in *NLRP3-/-* in **B**. GBM and **E**. astrocyte cells. **C,F**. MTT assay for cell viability in *NLRP3-/-* in GBM and astrocyte cells. Data are represented as mean ± SEM. Error bars indicated SEM. **H.** ELISA to determine cytokine levels of IL-6 (i), IL-1β (ii), TNF-⍺ (iii) in *NLRP3- /-* in GBM and astrocyte cells. Data are represented as mean ± SEM. Error bars indicated SEM. ∗p < 0.05, **p < 0.01, ***p < 0.005 (Student’s t-test).

Cytokine profiling of culture supernatants revealed that NLRP3 si-knockdown did not affect IL-1β secretion in LN-229 cells and enhanced IL-1β in SVG cells (Fig. 2G. IL-6 and TNF-⍺ secretion enhanced in LN-229 cells and suppressed in SVG cells, upon NLRP3 si-knockdown (Fig. 2H-I). These findings suggest that NLRP3 supports proliferation, migration, and viability in GBM cells, in contrast to its inhibitory role in astrocytes, highlighting a cell context-dependent function in the brain tumor microenvironment, similar to NLRP12 (21).

### NLRP3 deficiency alters glioma spheroid shape and cytokine levels, affecting tumor growth

Topological factors like tumor circularity and compactness have been used as diagnostic and prognostic markers for endometrial cancer and invasive cervical cancer (28,29). To recapitulate tumor architecture, cellular heterogeneity, and microenvironment, three-dimensional (3D) multicellular spheroids were generated using LN-229 and SVG monoculture and heterocellular spheroids with or without NLRP3 (Fig. 3A, Supplementary data Fig. S1). Quantification of the circularity and compactness revealed the NLRP3-deficient LN-229 monoculture spheroids exhibited decreased circularity, and compactness leading to an overall increased area, whereas the wild-type spheroids showed circular and tight spheroids in 24hrs (Day 0-1)(Fig. 3B-E). Heterocellular spheroids with wild-type LN-229 and SVG cells demonstrated compact and circular spheroids from Day 2-3, whereas compromised morphology was prominent on NLRP3-deficient heterocellular spheroids. A similar effect was observed with NLRP3 knockdown in either of the cells in the heterocellular spheroid environment. 24-hour temporal videos of spheroid formation were observed (Supplementary data Movie 1). Cytokine analysis revealed increased secretion of pro-inflammatory cytokines, including IL-1β, IL-6, and TNF-α, upon NLRP3 knockdown in both LN-229 and SVG monoculture spheroids. A similar trend has been observed in NLRP3 knockdown in either or both cells of a heterocellular spheroid, compared to wild-type cells (Fig. 3F-H). This suggests a pro-inflammatory tumor microenvironment that may support tumour growth.

**Fig 3.**
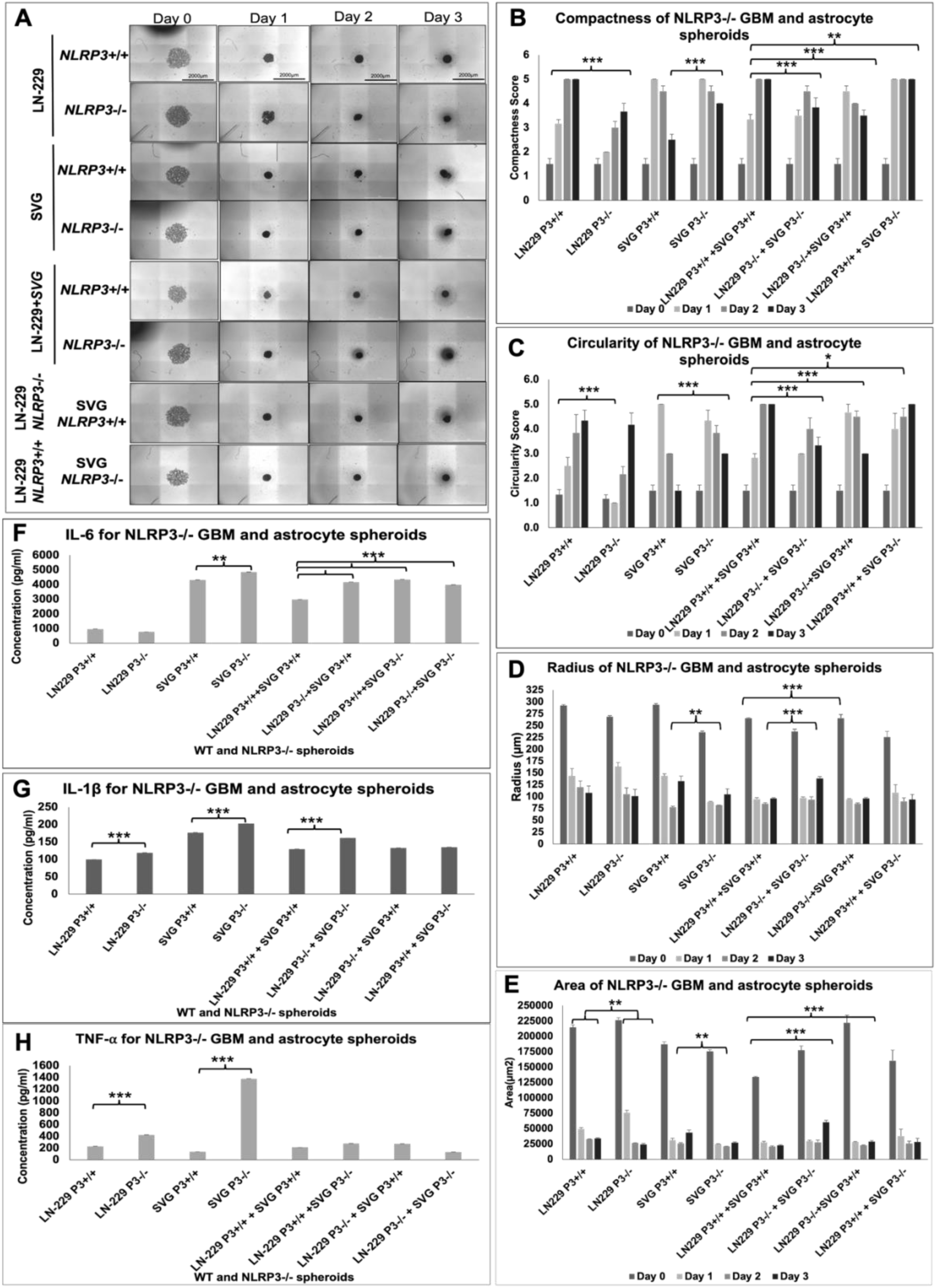
*NLRP3* si-knockdown in LN-229 and SVG affects spheroid circularity and compactness. **A**. Imaging of co-cultured spheroid for 3 consecutive days. **B**. circularity quantification (n=3). **C.** compactness quantification **D.** area quantification (n=3). ELISA to determine cytokine levels of **F.** IL-1β, **G**. IL-6, **H**. TNF-⍺ in *NLRP3* si-knockdown LN-229:SVG co-culture spheroids. Data are represented as mean ± SEM. Error bars indicated SEM. ∗p < 0.05, **p < 0.01, ***p < 0.005 (Student’s t-test).

### NLRP3 is Differentially expressed in patient-derived GBM tissues and organoids

Patient-derived tissues are widely used for diagnosis and expression analysis. Mouse models have traditionally been employed as a standard experimental system for investigating mechanisms relevant to human physiology (30,31). Patient-derived organoid models offer a clinically relevant platform for investigating intra-tumoral heterogeneity and tumor biology within the complex GBM tumor microenvironment (32–34). To comprehensively characterize NLRP3 expression across cellular heterogeneity and tumor grades, we analyzed primary glioma tissues and corresponding adjacent normal samples, as well as patient-derived three-dimensional organoid cultures. Patient-derived glioma and matched normal tissues were obtained from AIIMS Jodhpur (Fig. 5, Table 2). Fresh specimens were enzymatically dissociated using a standardized protocol (22) to obtain a heterogeneous population of glial cells. These cells were subsequently cultured to generate three-dimensional organoids over a period of seven days. Due to an ongoing patent application, specific details of the organoid generation protocol are confidential and cannot be disclosed. Total RNA and protein were extracted from matched tissue samples for molecular analyses.

To analyse expression fold changes of glioma-associated genes, the patient samples were quantified for expression fold changes for isocitrate dehydrogenase 1 (IDH1) wild-type (WT) and IDH1^R2H^, mutant, IDH2, epidermal growth factor receptor (EGFR), and tumor suppressor TP53 genes, by RT-qPCR, normalized against 18S rRNA and GAPDH controls (Supplemental data, Fig. S2) (2,20). *IDH1* and *IDH2* mutations, particularly the *IDH1* R132H substitution, represent hallmark genetic events in grade II and III gliomas, modulating tumor metabolism through aberrant accumulation of the oncometabolite R(–)-2-hydroxyglutarate (R-2HG), which influences cell cycle regulation and tumorigenic potential(36). In contrast, alterations in *TP53* and *EGFR* signaling pathways serve as principal drivers of glioblastoma pathogenesis, contributing to dysregulated cell-cycle progression, impaired apoptotic control, and enhanced proliferative signaling (35,36).

Table 2. Lists 12 patient sample details used in this study for expression analysis of NLRP3. The samples obtained were of different grades, two Grade 2, three 3, three Grade 4 and four Grade 4 samples (P08, P09, P10, and P11) were used for organoid generation. Three paired grade 3 tumors (Glioma, G), and tumor adjacent tissue (Normal, N) (P02, P03, and P04) expressed distinct NLRP3 expression patterns. P01, a grade 2 glioma patient, displayed high NLRP3 expression in the western blot assay (Fig. 4C i-ii). Grade 3 patient pairs expressed variable expression patterns. P02 exhibited high NLRP3 transcriptional expression, with more than a 1-fold change in the expression of IDH^R2H^, IDH2, EGFR, and TP53 genes (Fig. 4A-B). Western blot analysis shows low expression at the translational level (Fig. 4C i-ii). P03 demonstrated increased NLRP3 expression in glioma tissue as compared to its adjacent normal tissue, with more than a 1-fold change of gene expression for IDH^R2H^, IDH2, EGFR, TP53, and NLRP3 genes (Fig. 4A-B). P04 showed elevated expression of all the genes with more than 50-fold change at quantitative PCR (Fig. 4B) and higher NLRP3 in glioma as compared to normal using end-point PCR (Fig. 4A). Western blot analysis revealed elevated NLRP3 expression in both glioma and adjacent non-tumorous tissues, with comparatively higher levels observed in glioma samples (Fig. 4C i-ii). P05, a grade 2 glioma, showed high NLRP3 expression in western blot (Fig. 4C i-ii), and high fold change in IDH1-WT, IDH^R2H^, and EGFR expression (Fig. 4H). P06, a grade 4 GBM, exhibited high expression of NLRP3 and all the marker genes with more than 1 fold change in all the genes (Fig. 4H). Similarly, P07 and P12, both grade 4 GBM, highly expressed NLRP3. P07 showed a high fold change in IDH^R2H^, whereas P12 showed high IDH1 WT and IDH^R2H^ expression, along with IDH2, EGFR, and TP53 (Fig. 4H).

**Fig 4.**
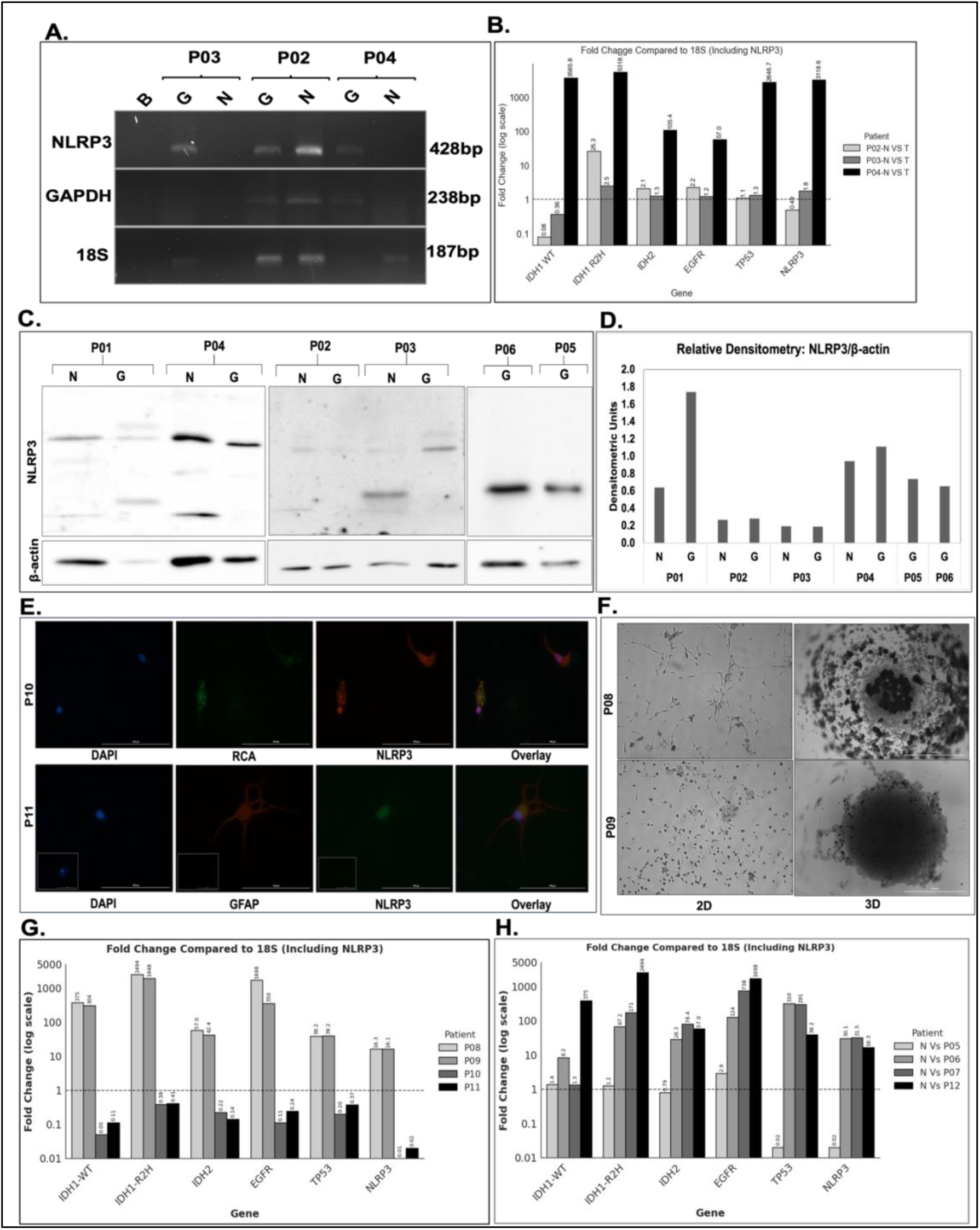
**Differential expression of Patient-derived Tissue**. List of patient-derived samples (Table 2). **A.** Agarose gel for Grade 3 Normal and Tumor Pair samples. Real-Time PCR to determine fold change in Glioma marker genes in patient-derived samples. **B**. Grade 3 Normal and Tumor Pair samples, **C.** Western Blot for NLRP3 protein expression in a patient pair and a single patient-derived tissue protein. **D**. Densitometry for western blot assay. **E.** Immunofluorescence study to determine NLRP3 (GFP) in patient-derived cells. **F.** 2D patient-derived cells culture vs 3D patient-derived organoids. G. Real-time PCR for Grade 4 tumor organoids. H. Real-time PCR for Grade 2 and Grade 4 Tumor samples. Data are represented as mean ± SD. Error bars indicated SD. ∗p < 0.05, **p < 0.01, ***p < 0.005 (Student’s t-test).

**Fig 5.**
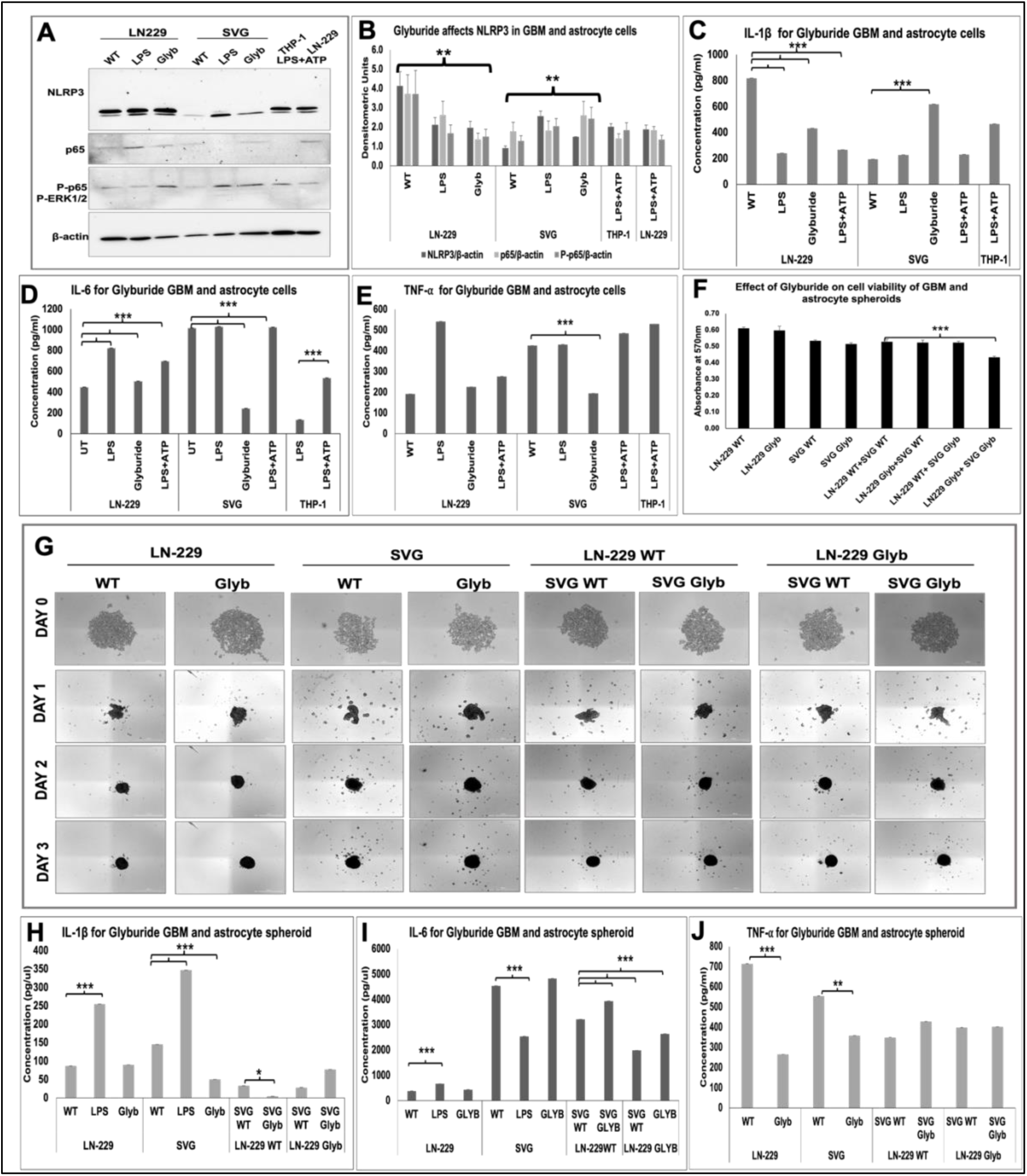
**Glyburide decreases NLRP3 expression in GBM Cell line**: **A**. Western Blot for UT, Glyburide and LPS-treated GBM and astrocyte cells (n=2) **B**. Densitometric analysis normalized to β-actin. **C.** ELISA to determine cytokine levels of C. IL-1β, D. IL-6, E. TNF-⍺ in Glyburide-treated cells. F. MTT Assay to access cell viability in WT and Glyburide-treated LN-229:SVG co-culture spheroids (n=2). G. WT and Glyburide treated monoculture and co-culture. ELISA to determine cytokine levels of H. IL-1β, I. IL-6, J. TNF-⍺ in WT and Glyburide-treated LN-229:SVG co-culture spheroids. Data are represented as mean ± SEM. Error bars indicated SEM. ∗p < 0.05, **p < 0.01, ***p < 0.005 (Student’s t-test).

Surgically resected Grade 4 tumor tissues (P08, P09, P10, and P11) were processed for 3D organoid culture and analyzed for 7 days. Immunofluorescence revelaed robust NLRP3 expression in P10 (Fig. 4E) and P08, P09 (Supplementary data). P10 cells with RCA staining and P11 for GFAP showed NLRP3 expression in microglia and astrocyte cells isolated from the patient tissue (Fig. 4E). Fig. 4F showed 2D isolated cells from tumor tissue and 3D organoids on day 3 after seeding. RNA isolated from organoids on day 7, was used for quantification using quantitative PCR. P08 and P09 showed a high fold change of genes IDH1-WT (>300), IDH^R2H^ (>1900), IDH2 (>40), EGFR(>350), TP53 (>35), and NLRP3 (>16) (Fig. 4G). P10 and P11 showed low expression of all the genes.

### Glyburide decreases NLRP3 expression in GBM and increases in astrocyte cells

Glyburide, a sulfonylurea class of antidiabetic drug, is a widely used drug for the treatment of Type 2 diabetes (37). The mechanistic pathway for glyburide is through high-affinity binding to the sulfonylurea receptor 1 (SUR1), the regulatory subunit of ATP-sensitive potassium (K^+^-ATP) channels in pancreatic β-cells. This binding inhibits channel activity, leading to membrane depolarization and the subsequent opening of voltage-gated calcium channels. The consequent elevation of intracellular calcium promotes insulin granule exocytosis, thereby stimulating insulin secretion (25). Glyburide has been reported to suppress NLRP3 inflammasome–mediated inflammation and associated colorectal tumorigenesis (38). Based on these findings, we sought to investigate whether Glyburide could similarly inhibit NLRP3 activation in glioblastoma and consequently attenuate tumorigenic progression. Glyburide-treated and WT LN-229 GBM and SVG astrocyte cells and 3D spheroids were investigated for NLRP3 expression. Western blot analysis reveals decreased NLRP3 expression in GBM cells, whereas increased NLRP3 expression in astrocyte cells as compared to wild-type (Fig. 5A i-ii). Cytokine profiling revealed a reduction in IL-1β secretion, accompanied by an increase in IL-6 levels, in LN-229 cells (Fig. 5B-C). In SVG astrocytic cells, IL-6 expression was similarly increased, whereas TNF-α levels were reduced (Fig. 5C-D). No significant alteration in TNF-α expression was observed in LN-229 cells (Fig. 5D).

Monocultured LN-229 and SVG, along with co-cultured spheroids, were seeded with and without Glyburide treatment to check for circularity, compactness, area, and radius of the tumor formation. No significant difference was observed in the tumour morphology. Whereas reduced cell viability was observed in co-cultured glyburide-treated LN-229: SVG spheroid as compared to wild-type (Fig. 5.5 F). Cytokine profiling demonstrated a reduction in IL-1β and TNF-α levels, in LN-229 and SVG cells (Fig. 5.5 C, 5.5 D, and 5.5 E). Glyburide treatment showed reduced cell viability in co-culture spheroids as compared to untreated spheroids (Fig. 5.5 F). No significant alteration in IL-6 expression was observed in LN-229 and SVG monocultured spheroids (Fig. 5.5 I). Co-cultured spheroids showed a significant reduction in IL-6 levels upon glyburide treatment, but no significant change for IL-1β and TNF-α was observed (Fig. 5.5 H-J).

## Discussion

Nucleotide-binding domain leucine-rich repeat-containing receptors (NLRs) act as intracellular sensors that control inflammation and cell death by forming inflammasomes and regulating cytokine release (39). Recent studies show that different NLR family members have context-specific roles in cancer, functioning as either tumor promoters or suppressors depending on the tumor type and microenvironment (40,41). Although the tumor microenvironment (TME), recurrence mechanisms, and molecular alterations in glioblastoma (GBM) have been extensively investigated, the specific functions of NLRs within glioma and GBM cells remain poorly defined ((3,9,42). According to recent reports, NLRP3 promotes glioma progression by activating the NF-κB signaling pathway (14). Our earlier work identified differential expression and a significant inverse correlation between methylated CpG loci and expression in GBM, for *NLRP3* and other NLR genes by using TCGA-GBM and low grade glioma (LGG) patient details (21). In this study, we further explored the NLRP3 expression using GBM and astrocyte (normal) cell lines, patient-derived tissue and patient-derived organoid models. Although NLRP3 has been reported to be highly expressed in U87 and U251 cell lines, we report NLRP3 expression in LN-18, LN-229 GBM, and SVG astrocyte cells. Basal expression of NLRP3 was found to be higher in astrocytes through immunocytochemistry, accompanied by noticeable alterations in astrocytic morphology following LPS stimulation. Additionally, increased punctate cytoplasmic localization of NLRP3 surrounding the nucleus was observed, suggesting inflammasome activation dynamics in response to inflammatory signaling. (Fig. 1C). However, no morphology change was observed for LN-229 GBM cells. Both transcriptional and translational analyses revealed an upregulation of NLRP3 expression in LN-229 and SVG cells following LPS stimulation (Fig. 1A, B, and D).

Functionally, siRNA-mediated silencing of NLRP3 resulted in decreased proliferation, migration, and viability in LN-229 cells, consistent with previous observations in other GBM models (14). In contrast, this study provides the first evidence that NLRP3 si-knockdown in non-neoplastic astrocytes markedly enhances their proliferation, viability, and migratory capacity (Fig. 2). NLRP3-deficient astrocytes exhibited increased IL-1β but reduced IL-6 and TNF-α secretion, suggesting a shift toward a paracrine, tumor-supportive phenotype within the GBM microenvironment. Conversely, NLRP3-deficient GBM cells showed elevated IL-6 and TNF-α with unchanged IL-1β levels, reflecting sustained pro-inflammatory signaling that may promote crosstalk with neighbouring cells and reinforce a pro-tumorigenic niche (Fig. 2G–I).

Considering the pronounced intra- and inter-tumoral heterogeneity that characterizes GBM (1,3,31,33,43,44), heterocellular spheroids, patient-derived tissues, and organoid models were employed to investigate NLRP3 expression patterns within a clinically relevant framework. NLRP3-deficient GBM monoculture spheroids showed compromised organizational integrity and cell-to-cell adhesion, resulting in disrupted spheroid compactness and circularity(Fig. 3). This aligns with reports stating NLRP3 knockdown decrease cell proliferation, invasiveness and migration (14). We observed differential expression of NLRP3 across different tumor grades, with consistently high NLRP3 expression in the tumor as compared to the tumor-adjacent normal sample (Fig. 4).

Multiple drugs targeting NLRP3 and its downstream pathways are in clinical trials to fight various autoimmune diseases (37,45,46). MCC950 is a small molecule inhibitor of NLRP3 activation and downstream IL-1β production in mouse models for Cryopyrin-associated periodic syndrome (CAPS), Multiple sclerosis, and Muckel-Wells syndrome(47,48). GDC-2394 is another small molecule inhibitor of NLRP3 activation in disease conditions like periodic fever, coronary artery disease (CAD), gout, etc. First-phase clinical trials in humans for CAD showed improved conditions, with reduced NLRP3 activation and decreased production of IL-1β and IL-18 (46). However, these studies require further research on the extent of inhibition for clinical efficiency in patients(46). Glyburide is one such anti-diabetic drug that inhibits NLRP3 inflammasome activation (25). In this study, glyburide treatment led to a reduction in NLRP3 expression, accompanied by decreased secretion of IL-1β, IL-6, and TNF-α (Fig. 5C–E), suggesting its potential therapeutic value in GBM management. However, in astrocytes, glyburide treatment resulted in elevated NLRP3 expression and increased IL-1β release, indicating a possible risk of inflammation arising from tumor-adjacent normal cells.

In summary, our data reveal that NLRP3 expression was elevated in GBM compared to adjacent normal tissue and increased further upon LPS stimulation. Functional assays revealed that NLRP3 knockdown reduced proliferation, viability, and migration in GBM cells but enhanced these processes in astrocytes, accompanied by distinct cytokine secretion profiles. These findings highlight the dual, context-dependent role of NLRP3 in tumor and non-tumor cells within the GBM microenvironment. While glyburide curbs GBM inflammation, astrocyte activation warrants the use of selective inhibitors or a combination with anti-IL-1β. Future studies aimed at elucidating NLRP3-mediated signaling mechanisms and their crosstalk with the tumor microenvironment may provide critical insights for the development of novel, targeted therapeutic strategies against this aggressive malignancy.

## Ethics Approval and Consent to Participate

Glioma tissues were obtained with approval from the Internal Review Board and the Ethics Committees of AIIMS, Jodhpur. Informed consent was acquired from human participants for the use of tissue samples for experiments. All experiments were performed in accordance with the ethical guidelines and regulations of AIIMS, Jodhpur, and Indian Institute of Technology Jodhpur.

## Competing Interests

The authors declare to have no competing interests that might be perceived to influence the results and/or discussion reported in this paper.

## Funding

This work had no external funding and was supported by institutional grants from the Indian Institute of Technology Jodhpur (IITJ), Rajasthan.

## Author Contribution

S.R. designed and performed the experiments (data analysis, cell culture, immunofluorescence, western blotting, primary cell isolation, RT-PCR, siRNA-mediated knockdown experiments, 3D spheroids, and patient-derived organoid generation, MTT, CFA, and migration assays, ELISA) and prepared and edited the initial manuscript draft. L.S. helped with ELISA experiments. D.M. helped with cell culture. D.K quantified cell and colony numbers, the spheroid’s circularity and compactness. R.M.S. developed the code for circularity and compactness. D.J., V.J., M.G., and M.A. provided patient tissue samples for ICC and western blot. S.J. conceptualized the study, designed experiments, and edited and reviewed the manuscript. All authors reviewed the manuscript.

## Supporting information

NLRP3 Supplementary Data

## Acknowledgement

This work was supported by institutional grants from the Indian Institute of Technology Jodhpur (IITJ) and funding from the Ministry of Electronics and Information Technology, Government of India (Grant No. 4(16)/2019-ITEA). We are grateful to Dr. Pankaj Seth (National Brain Research Centre, NBRC) for kindly providing the human astrocyte cell line SVG. We thank Mr. Bharat Pareek, Technical Superintendent at IIT Jodhpur, for his invaluable technical support in the laboratory. We also acknowledge the critical contributions of Devansh Shah (RT-PCR analyses), Kane Chaitravi Anand (quantification of colony formation assay), and Manogna Swayampakula, Mukesh Kumar, Hemant Sunaliya, and Rajdip Panda Mahapatra (quantification of spheroid circularity, compactness, and area). Additionally, we appreciate the efforts of Pooja Porwal, Nidhi Patel, Aditya Parkhi, Krishna Dev Solanki, Rohitansh Bishnoi, Mayank Srivastava, and Piyush Khandari for their assistance with cell migration quantification. These contributions were part of their B.Tech. research projects.

## Notes

### Competing Interest Statement

The authors have declared no competing interest.

