## Supplementary material for "NLRP3 Inflammasome Exhibits Context-Dependent Roles in Glioblastoma-Astrocyte Crosstalk": NLRP3 Supplementary Data

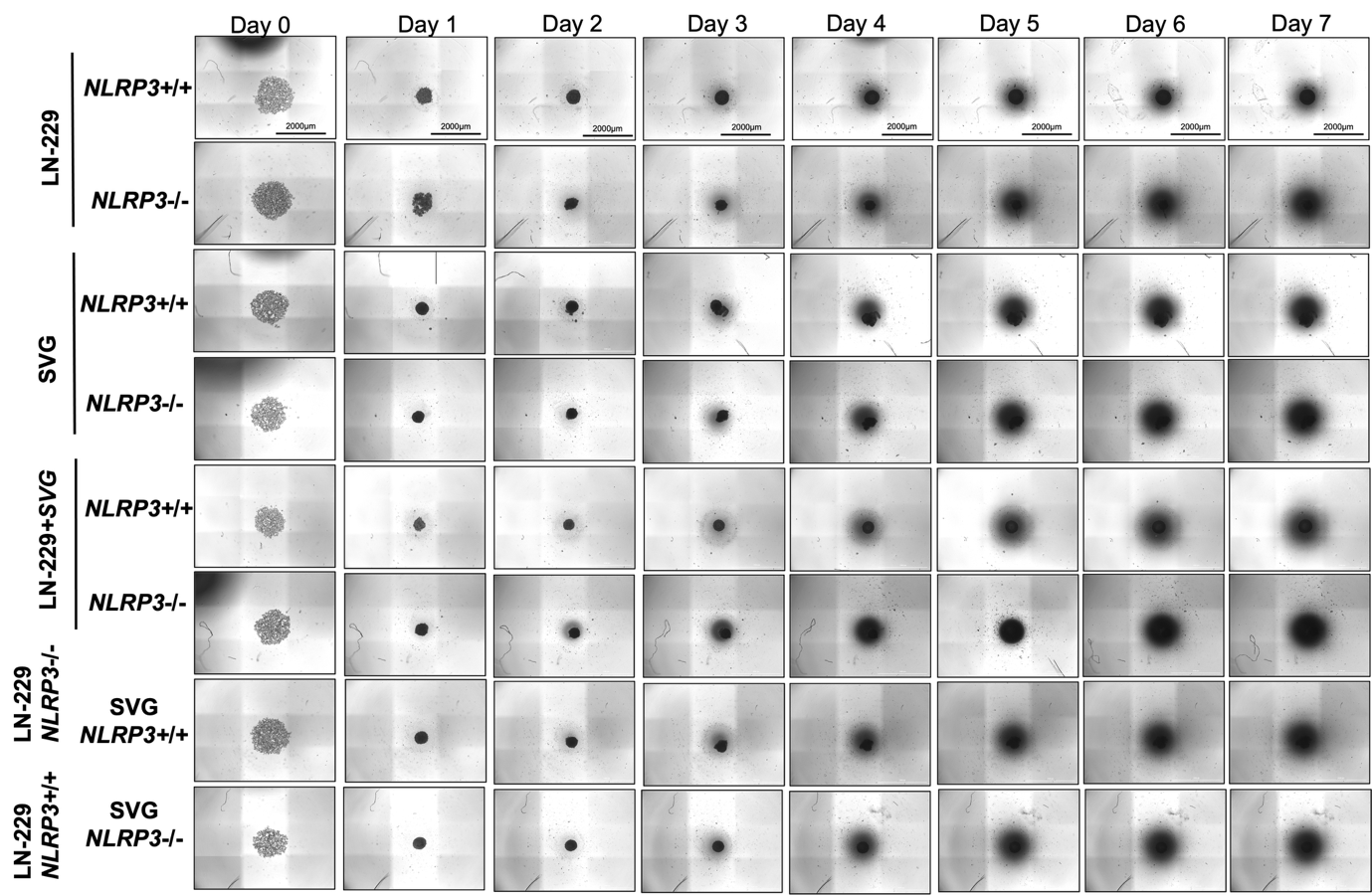

Figure S1**. *NLRP3* si-knockdown in LN-229 and SVG affects spheroid circularity and compactness.** **A**. i) Imaging of co-cultured spheroid for 7 consecutive days. Supplementary to Figure 3.

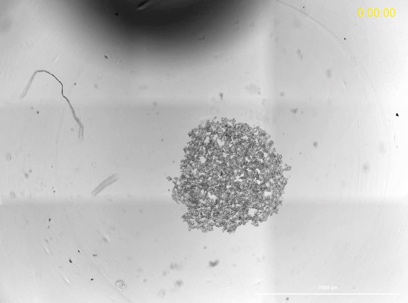

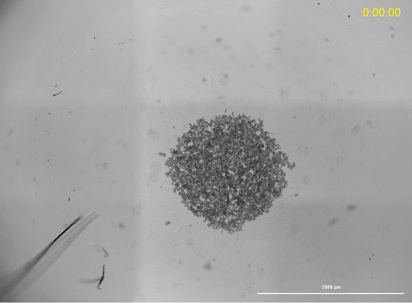

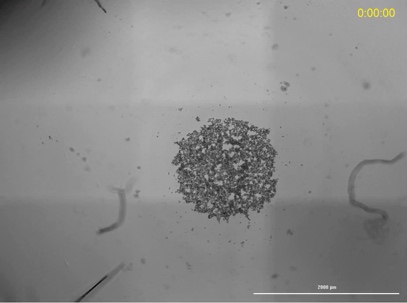

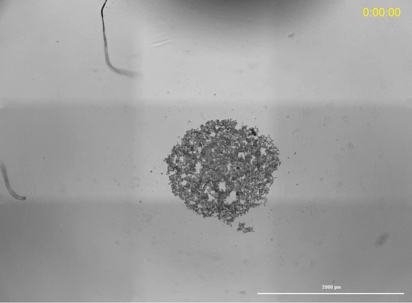
LN-229 WT	LN-229 NLRP3-/-

SVG WT	SVG NLRP3-/-

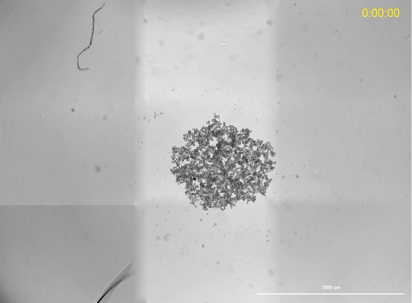

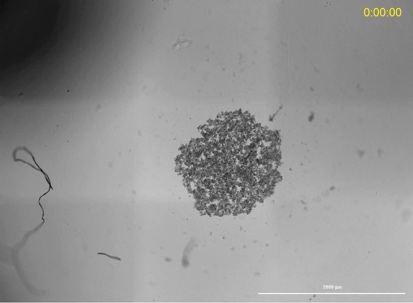

LN-229 WT+SVG WT	LN-229 P3-/-+SVG P3-/-

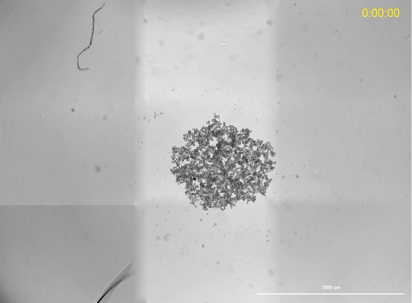

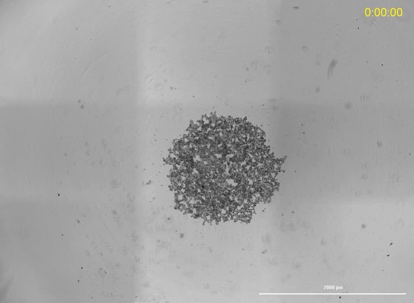

LN-229 P3-/-+SVG WT	LN-229 P3 WT+SVG P3-/-

**Movie 1. Process of spheroid generation, related to Figure 3:** Movies of LN-229 and SVG monoculture and co-culture spheroids. Cells were seeded for spheroid formation, and live cell imaging was performed for 48 hours. For the creation of the movie, images were taken at 30-minute intervals. Movies are representative of 3 experiments. Movies were taken at a 4X objective lens, Scale bar, 2000 μm.

A.

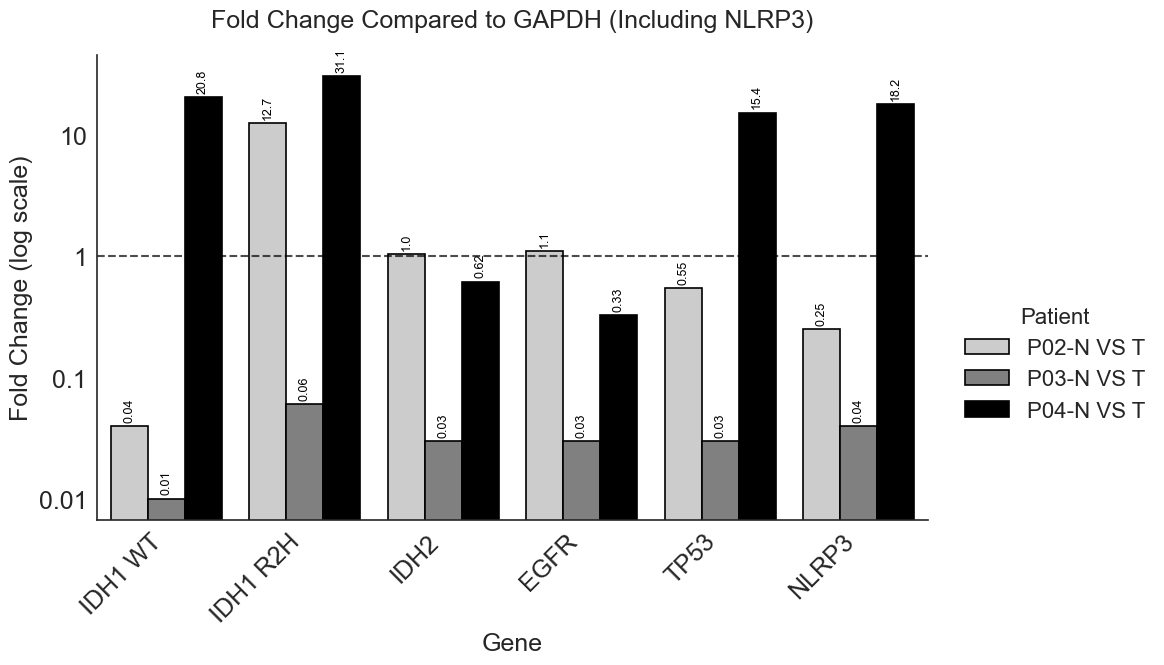

B.

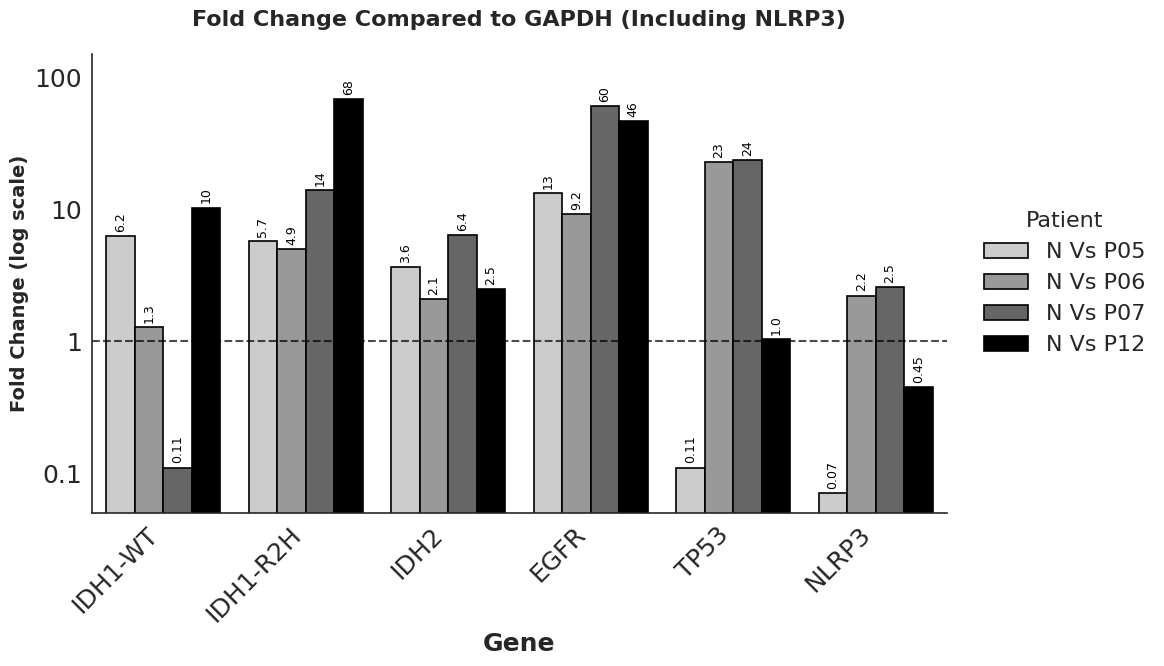

C.

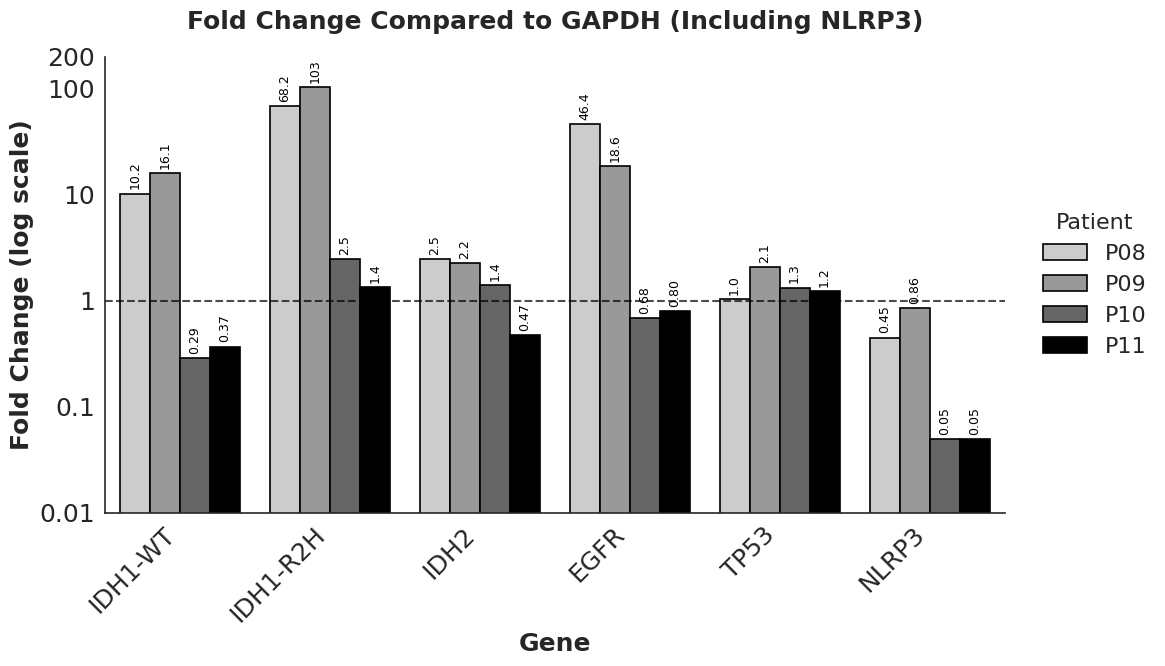

**Figure S2**. **Differential expression of Patient-derived Tissue**. List of patient-derived samples (Table 2). Real-Time PCR to determine fold change in Glioma marker genes in patient-derived samples. (A) Grade 3 Normal and Tumor Pair samples, (B) Grade 4 Tumor samples, (C) Grade 4 patient-derived organoids.
